# Effects of Copy Number Variations on Longevity in Late-Onset Alzheimer’s Disease Patients: Insights from a Causality Network Analysis

**DOI:** 10.1101/2023.07.04.547622

**Authors:** Yanan Hao, Chuhao Li, He Wang, Chen Ming

## Abstract

Alzheimer’s disease (AD), particularly late-onset Alzheimer’s disease (LOAD), is a prevalent form of dementia that significantly affects patients’ cognitive and behavioral capacities and longevity. Although approximately 70 genetic risk factors linked with AD have been identified, their influence on patient longevity remains unclear. Further, recent studies have associated copy number variations (CNVs) with the longevity of healthy individuals and immune-related pathways in AD patients. This study aims to investigate the role of CNVs on the longevity of AD patients by integrating multi-omics data from the Religious Orders Study/Memory and Aging Project (ROSMAP) cohort through causality network inference.

Our comprehensive analysis led to the construction of a CNV-gene-age of death (AOD) causality network. We successfully identified three key CNVs (DEL5006, mCNV14192, and DUP42180) and seven AD-longevity causal genes (*PLGRKT, TLR1, PLAU, CALB2, SYTL2, OTOF,* and *NT5DC1*) impacting AD patient longevity, independent of disease severity. This outcome emphasizes the potential role of plasminogen activation and chemotaxis in longevity. We propose several hypotheses regarding the role of identified CNVs and the plasminogen system on patient longevity.

However, experimental validation is required to further corroborate these findings and uncover precise mechanisms. Despite these limitations, our study offers promising insights into the genetic influence on AD patient longevity and contributes to paving the way for potential therapeutic interventions.

## 1 Introduction

Alzheimer’s disease (AD), the most prevalent form of dementia, is estimated to affect approximately 40 million individuals globally based on estimations from WHO (Reitz *et al*. 2023; Alzheimer’s Disease International (ADI)). The late-onset Alzheimer’s disease (LOAD), diagnosed typically after the age of 65, accounts for about 95% of all AD cases (Bekris *et al*. 2010; Bertram & Tanzi 2005). Characterized by beta-amyloid plaque and neurofibrillary tangles (DeTure & Dickson 2019), LOAD represents a profound impact on not only cognitive and behavioral capabilities but also patient longevity. After clinical diagnosis, the average life expectancy of AD patients ranges from four to eight years, though some cases report living up to 20 years with the disease (Alzheimer’s Association). Notably, there exists a significant variation in life expectancy among AD patients, and the underlying genetic mechanisms contributing to this variability remain unknown.

Large-scale cohort studies focusing on Alzheimer’s disease have identified approximately 70 genetic risk factors linked with AD (Reitz *et al*. 2023; Marioni *et al*. 2018; Lambert *et al*. 2013; Kunkle *et al*. 2019; Jansen *et al*. 2019; Wightman *et al*. 2021; Bellenguez *et al*. 2022). Although these factors may contribute to the predisposition to AD, their impact on the longevity of patients remains largely unexplored. A previous study based on 2,872 Danish twins has observed that genetic factors can explain 25% variation in the human lifespan (Herskind *et al*. 1996). And several previous research has shed light on the mechanisms by which genes contribute to longevity (Conneely *et al*. 2012; Melzer *et al*. 2020). Meanwhile, several Danish and U.S. population-based cohort studies have found that genome-wide copy number variation (CNV) burden is associated with human longevity (Nygaard *et al*. 2016; Kuningas *et al*. 2011). A Han Chinese population-based GWAS study also discovered that CNVs are associated with human longevity through various aging-related phenotypes, such as telomere length, the risk of cancer, and vascular and immune-related diseases (Zhao et al., 2018). Intriguingly, our previous study revealed an association between CNVs and immune-related pathways in AD patients through multi-omics integration (Ming *et al*. 2022).

Motivated to unravel the influence of CNVs on the longevity of AD patients, we conducted a comprehensive analysis integrating multi-omics data from the Religious Orders Study/Memory and Aging Project (ROSMAP) cohort (De Jager *et al*. 2018; Mostafavi *et al*. 2018a; Chibnik *et al*. 2018; Lee *et al*. 2023) (**Figure 1**). Our research focuses on elucidating the causality network connecting copy number variations, gene expression, and the age of death in Alzheimer’s patients. Through our comprehensive analysis, we identified three key CNVs (i.e., DEL5006, mCNV14192, and DUP42180) that regulate the expression of seven genes (i.e., *PLGRKT, TLR1, PLAU, CALB2, SYTL2, OTOF,* and *NT5DC1*) ultimately impacting the age of death of AD patients. The AOD-correlated genes further showed functional enrichment on the plasminogen activation and chemotaxis pathway. This research sheds light on the genetic factors contributing to the discrepancy in longevity observed among AD patients and provides insights into potential therapeutic targets for improving the longevity of AD patients.

**Figure 1.**
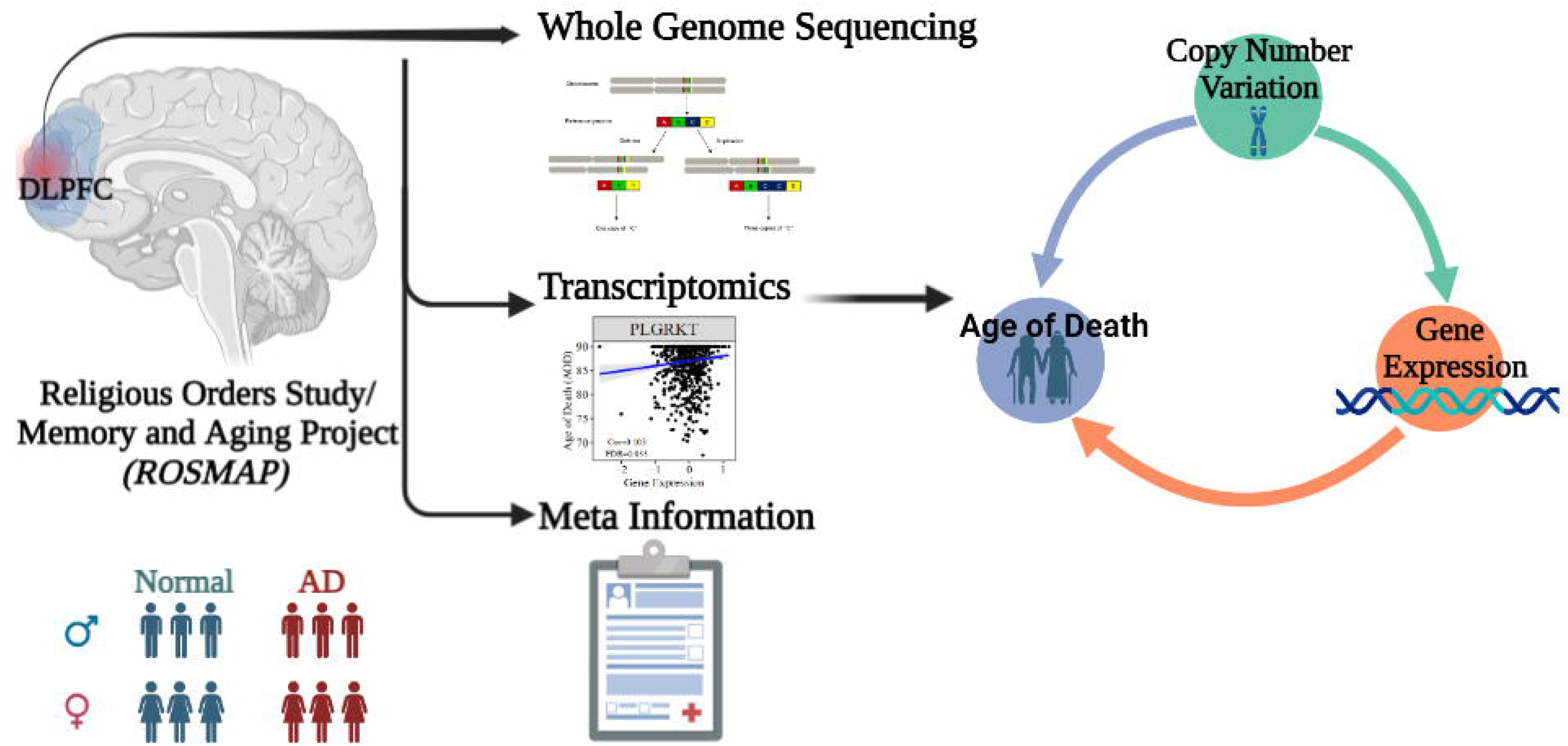
Abstract graph of integrating WGS-based CNV, transcriptomics data and meta information of ROSMAP cohort.

## 2 Results

### 2.1 Copy number dosages of ten CNVs are significantly correlated with the age of death of Alzheimer’s patients

We constructed a generalized linear regression model to investigate the correlation between CNVs and the age of death (AOD) of AD patients based on copy number profiles and meta-information of 1,127 North American White individuals from the ROSMAP AD cohort (De Jager *et al*. 2018; Mostafavi *et al*. 2018a; Chibnik *et al*. 2018; Lee *et al*. 2023) (**Table S1**), while sex and disease status were treated as covariances (**Methods 4.5, Figure 2**). There are 762 AD patients and 365 NL patients based on the physician’s overall cognitive diagnostic category at the time of death (cogdx) in the ROSMAP cohort. To exclude the bias from the severity of AD status, we treated various AD pathological traits, which are cogdx, Braak stage score (braaksc), and Consortium to Establish a Registry for Alzheimer’s Disease score (ceradsc), to represent the severity of AD independently in the generalized linear regression analysis (**Methods 4.5**). Upon accounting for potential biases linked to sex and disease status, our analysis revealed 12, 12, and 11 AOD-correlated CNVs when incorporating cogdx, braaksc, and ceradsc as covariates, respectively. These findings each maintained a false discovery rate (FDR) of less than 0.05(**Figure S1**). There were ten consensus AOD-correlated CNVs under all three AD pathological criteria in total (**Table 1**, **Figure 3**). Among them, six CNVs were positively correlated with AOD, while four CNVs were negatively correlated with AOD (**Table 1** and **Figure 3**).

**Figure 2.**
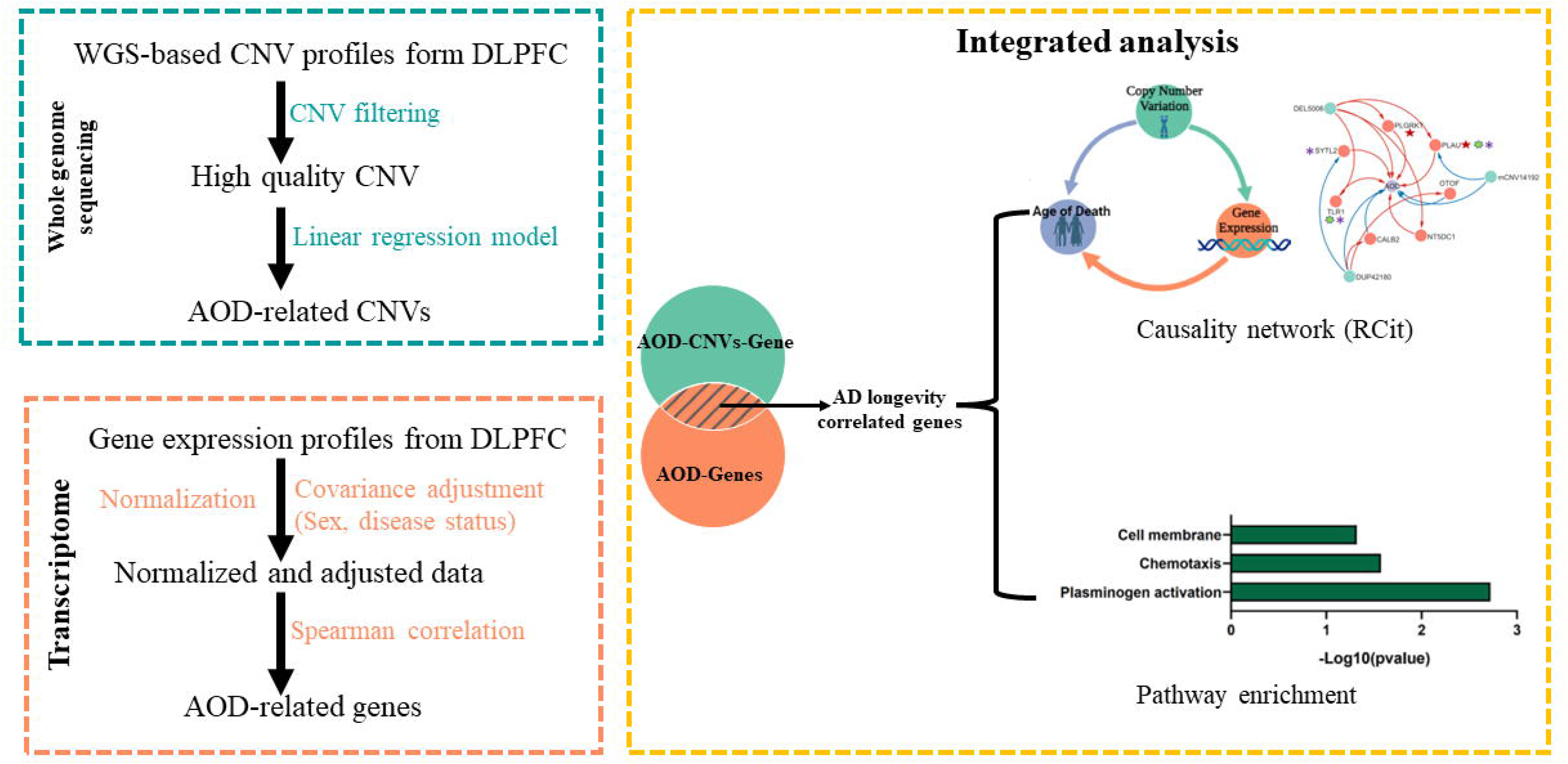
The pipeline to integrate WGS-based CNV and transcriptomics data. Green block, WGS-based CNV data; Orange block, transcriptomics data; Yellow block, integrated analysis for CNV and gene.

**Figure 3.**
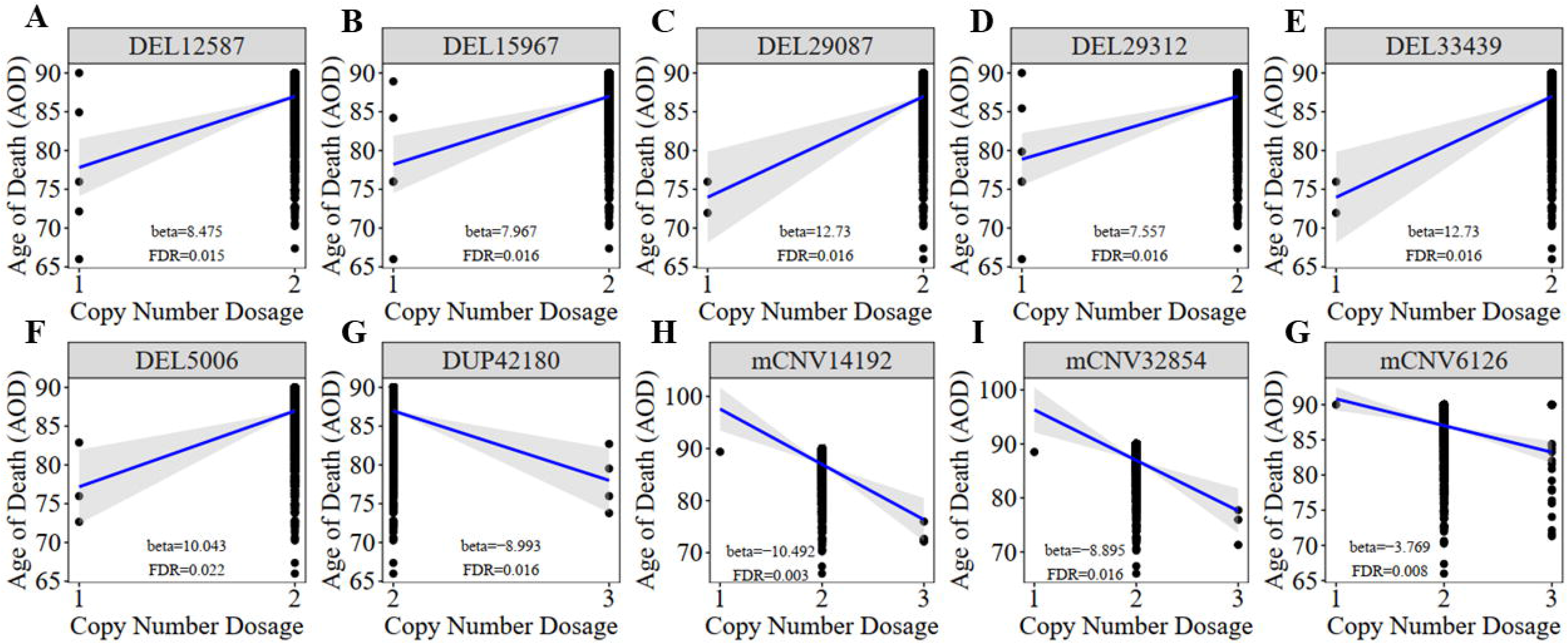
The correlation between AOD-correlated CNV and AOD. We utilized a generalized linear model in Matrix eQTL to perform correlation analysis between AOD-correlated CNV dosage and gene expression. t-statistic, False Discovery Rate (FDR) < 0.05; Beta: estimated effect size of CNV on AOD; Blue line: regression line added by R function ggscatter; Grey: the 95% confidence interval.

**Table 1.** Summary of the significant AOD-correlated CNVs.

| CNV | Chr | Start | End | Beta<br>(CNV-AOD) | P.Adj<br>(CNV-AOD) | CNV frequency<br>(ROSMAP) |
| --- | --- | --- | --- | --- | --- | --- |
| DEL33439 | chr1 | 28,670,388 | 28,698,217 | 1.27E+01 | 1.58E-02 | 0.18% |
| DEL12587 | chr3 | 103,265,515 | 103,266,967 | 8.48E+00 | 1.47E-02 | 0.44% |
| mCNV32854 | chr1 | 125,318,955 | 125,323,070 | -8.89E+00 | 1.58E-02 | 0.35% |
| DEL29312 | chr9 | 24,099,045 | 24,104,914 | 7.56E+00 | 1.58E-02 | 0.53% |
| mCNV14192 | chr3 | 171,925,368 | 171,927,349 | -1.05E+01 | 3.25E-03 | 0.35% |
| DUP42180 | chr1 | 31,476,591 | 31,489,411 | -8.99E+00 | 1.58E-02 | 0.35% |
| DEL5006 | chr2 | 17,194,382 | 17,196,437 | 1.00E+01 | 2.19E-02 | 0.27% |
| mCNV6126 | chr2 | 71,904,772 | 71,907,076 | -3.77E+00 | 7.97E-03 | 2.48% |
| DEL29087 | chr9 | 11,470,585 | 11,635,259 | 1.27E+01 | 1.58E-02 | 0.18% |
| DEL15967 | chr4 | 64,478,325 | 64,483,686 | 7.97E+00 | 1.58E-02 | 0.44% |

Note: Beta, estimated effect size; P.adj, adjusted P-value calculated based on linear regression by Matrix eQTL and adjusted by FDR; CNV frequency, the frequency of CNV in the ROSMAP cohort.

To investigate whether there are significant differences in frequency between the AD and healthy control groups for the above ten AOD-CNVs, we compared their frequency using the chi-square test (**Methods 4.6**). We found all 10 AOD-correlated CNVs with low frequency in both AD and healthy control groups, and they showed no significant frequency difference in both groups (**Table S2**). To further investigate the prevalence of the above AOD-correlated CNVs in the human population, we compared our AOD-correlated CNVs with the public CNV datasets GnomAD (Chen *et al*. 2022) based on large populations. 7 CNVs were validated in the public CNV databases and also showed rare frequency in human populations (**Table S2**).

### 2.2 The AOD-correlated CNVs regulate the expression of seventeen AOD-correlated genes in the DLPFC region of AD patients

To further explore how AOD-correlated CNVs regulate gene expression in the brains of AD patients, we implemented the expression quantitative loci (eQTL) analysis by using Matrix eQTL software (Shabalin 2012) to integrate genome-wide CNV profiles and transcriptomics data of the dorsal lateral prefrontal cortex (DLPFC) region from the ROSMAP cohort (**Methods 4.6**, **Figure 2**). Based on the linear additive model with sex and disease status as covariates, there were 110 genes identified with their expression level significantly correlated with copy number dosage of the 10 AOD-correlated CNVs under the FDR threshold at 5%, forming 125 CNV-Gene pairs (**Table S3**).

To pinpoint AOD-correlated genes, we further performed Spearman correlation analysis between all genes identified in the transcriptomics data of the DLPFC region and AOD (**Methods 4.7**). The transcriptomic profile was further adjusted for AD pathological traits using limma R package (Ritchie *et al*. 2015) to exclude bias from disease status (**Figure S2, Methods 4.7**). There were 2,056, 1,294, and 2,774 AOD-correlated genes identified based on cogdx, braaksc, and ceradsc respectively, under the genome-wide FDR threshold of 0.05 (**Table S4)**.

By intersecting the 2,056, 1,294 and 2,274 AOD-correlated genes with the 125 AOD-CNV correlated genes, we finally pinpointed 17, 14, 21 genes which were both correlated with AOD and AOD-CNVs respectively, defined as presumptive AD longevity-correlated genes. We used 17 genes underlying cogdx for further analysis and these genes formed 23 CNV-Gene-AOD pairs with 9 AOD-correlated CNVs. We further constructed correlation network for above CNV-gene-AOD pairs. (**Figure 3**, **Table S5, Methods 4.8**).

### 2.3 Causality network between CNVs, Gene expression, and Age of death of AD patients pinpointed three key CNVs, seven key genes and showed enrichment on plasminogen activation pathway

To elucidate the inherent relationship between variations in gene expression levels and aging, we established a causality network that integrated AOD-correlated CNVs, presumptive AD-longevity-associated genes, and AOD by implementing the Causal Inference Test (CIT)(Millstein *et al*. 2016)(**Methods 4.8**). After excluding the aging caused variation in gene expression, there are three CNVs (i.e., DEL5006, mCNV14192, and DUP42180) regulating the expression level of seven genes (i.e., *PLGRKT, TLR1, PLAU, CALB2, SYTL2, OTOF,* and *NT5DC1*), and the expression level of these seven genes further regulates the age of death of AD patients, in the final causality network (**Figure 5** and **Table 2**). We named the seven genes in the above causality network as AD longevity-causal genes. The copy number dosage of DEL5006 are positively correlated with most of AD longevity positively-causal genes (i.e., *PLGRKT*, *TLR1*, *PLAU* and *NT5DC1*), and served as a key longevity CNV loci. Meanwhile, the copy number dosage of the other two CNVs, which are DUP42180 and mCNV14192, are negatively correlated with AOD through regulating the expression of genes *OTOF*, *SYTL2*, *CALB2,* and *PLAU*. Interestingly, we further found two AD-risk genes, *PLAU* and *TLR1*, which functions in the Aβ clearance and degradation process (Ertekin-Taner *et al*. 2005; Liu *et al*. 2012) were positively correlated with AOD of AD patients. The copy number dosage of DEL5006 demonstrates a positive correlation with the expression level of *PLAU*. Conversely, the dosage of mCNV14192 shows a negative correlation with *PLAU* expression. The multi-CNV correlation pattern indicates the importance of *PLAU* in AD longevity. Notably, in the transgenic alpha murine urokinase-type plasminogen activator (αMUPA) mouse model, *PLAU* had been reported that its overexpression in the brain may extend mouse longevity by limiting food consumption (Miskin & Masos 1997), which is consistent with our observation in AD patients (**Figure 5**).

**Figure 4.**
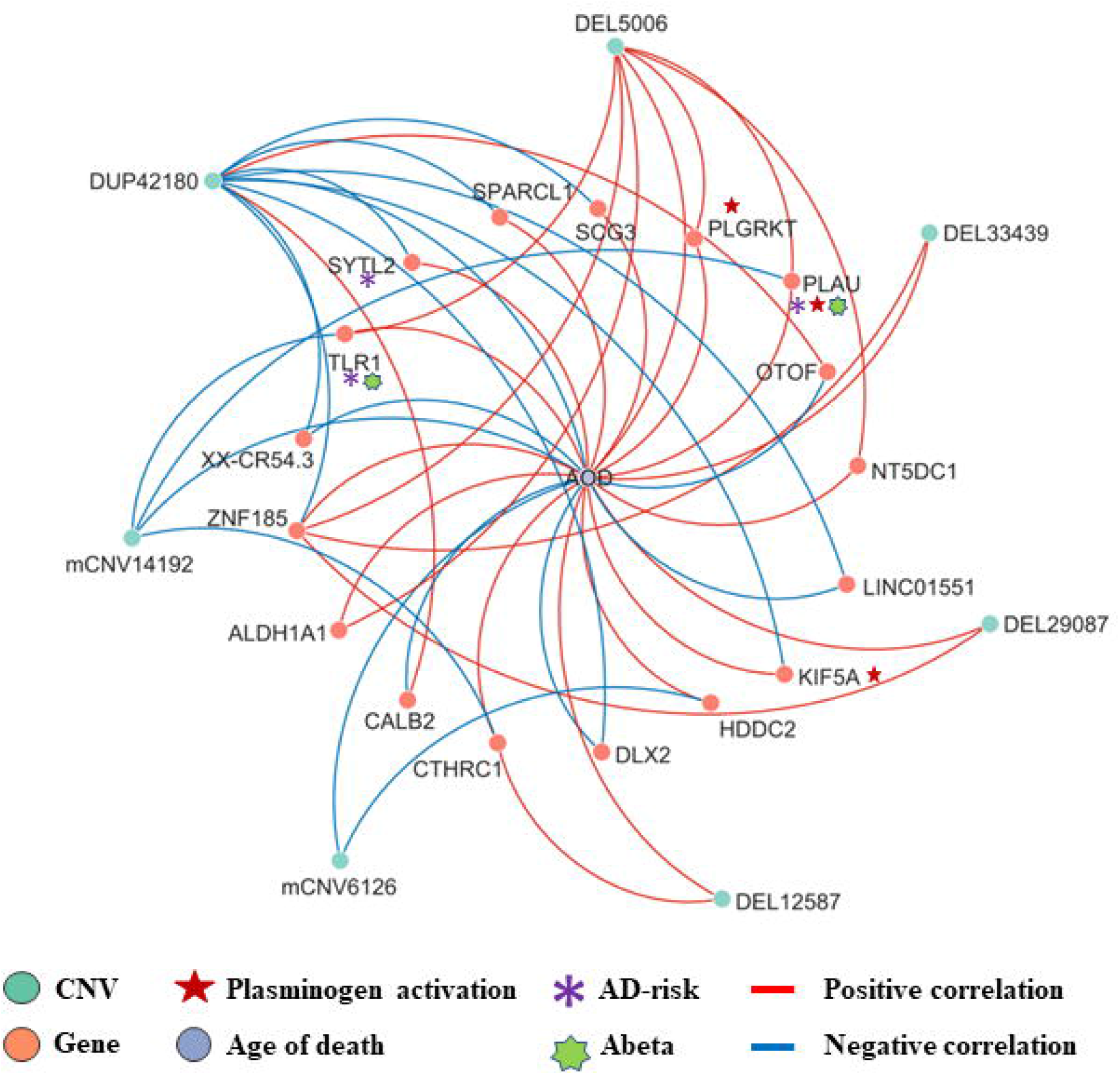
The correlation network between AOD-correlated CNVs, Gene expression, and AOD. The correlation network connected 9 AOD-correlated CNVs, 17 AOD-correlated genes, and age of death (AOD). Yellow: genes, Pink: AOD-correlated CNVs; Green: AOD; Red lines: positive correlation; Blue lines: negative correlation; Red five-pointed star: plasminogen activation-related genes; Green seven-pointed star, Abeta genes; Purple asterisk, AD-risk genes.

**Figure 5.**
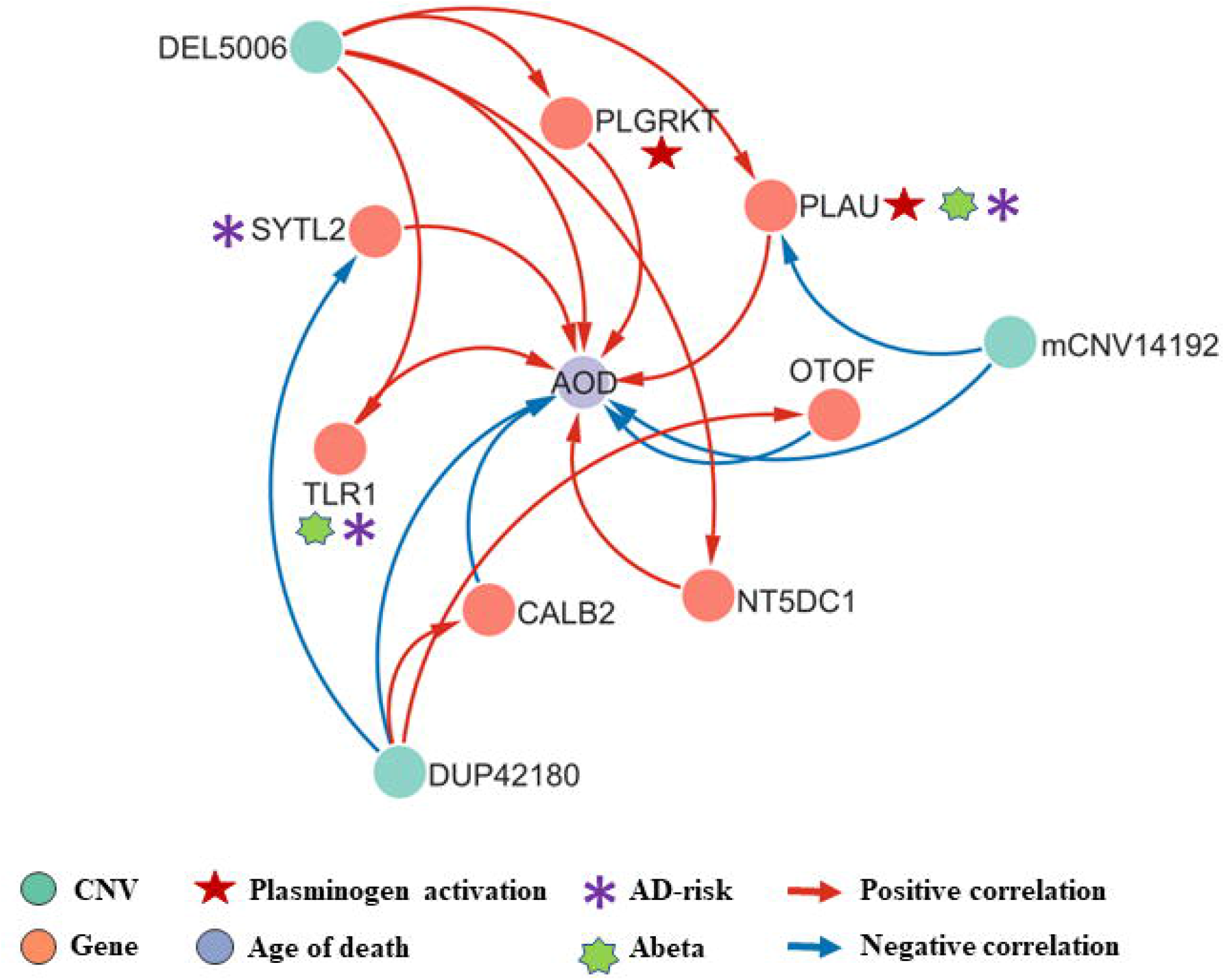
Causality network between AOD-correlated CNVs, AD longevity-causal genes, and AOD. The causality network connected 3 AOD-correlated CNVs, 7 AD longevity-causal genes, and AOD. Yellow: genes, Pink: AOD-correlated CNVs; Green: AOD; Red arrows: positive correlation; Blue arrows: negative correlation; Red five-pointed star: plasminogen activation-related genes; Green seven-pointed star, Abeta genes; Purple asterisk, AD-risk genes.

**Table 2.** The summary of CNV-gene-AOD pairs in the causality network based on cogdx.

| CNV | Gene | Trait | Beta<br>(CNV-Gene) | P.adj<br>(CNV-Gene) | Beta<br>(CNV-AOD) | P.adj<br>(CNV-AOD) | Cor.<br>coefficient<br>(Gene-AOD) | P.adj<br>(Gene-AOD) | P.adj_cit |
| --- | --- | --- | --- | --- | --- | --- | --- | --- | --- |
| DEL5006 | *PLGRKT* | AOD | 1.25E+00 | 1.71E-02 | 1.00E+01 | 2.19E-02 | 1.03E-01 | 3.27E-02 | 3.07E-02 |
| DUP42180 | *OTOF* | AOD | 1.15E+00 | 3.74E-02 | -8.99E+00 | 1.58E-02 | -1.65E-01 | 8.12E-04 | 3.85E-03 |
| mCNV14192 | *PLAU* | AOD | -1.52E+00 | 3.22E-03 | -1.05E+01 | 3.25E-03 | 1.33E-01 | 6.32E-03 | 3.81E-03 |
| DEL5006 | *PLAU* | AOD | 1.73E+00 | 4.28E-02 | 1.00E+01 | 2.19E-02 | 1.33E-01 | 6.32E-03 | 3.85E-03 |
| DUP42180 | *SYTL2* | AOD | -9.63E-01 | 1.96E-02 | -8.99E+00 | 1.58E-02 | 1.04E-01 | 3.18E-02 | 2.72E-02 |
| DUP42180 | *CALB2* | AOD | 1.77E+00 | 4.26E-02 | -8.99E+00 | 1.58E-02 | 1.35E-01 | 8.94E-03 | 1.28E-02 |
| DEL5006 | *TLR1* | AOD | 2.45E+00 | 2.67E-02 | 1.00E+01 | 2.19E-02 | 1.52E-01 | 5.45E-03 | 3.85E-03 |
| mCNV14192 | *TLR1* | AOD | -1.78E+00 | 2.04E-02 | -1.05E+01 | 3.25E-03 | 1.35E-01 | 5.45E-03 | 3.85E-03 |
| DEL5006 | *NT5DC1* | AOD | 9.95E-01 | 4.89E-02 | 1.00E+01 | 2.19E-02 | 1.52E-01 | 1.82E-03 | 3.85E-03 |
| DEL5006 | *PLGRKT* | AOD | 1.25E+00 | 1.71E-02 | 1.00E+01 | 2.19E-02 | 1.03E-01 | 3.27E-02 | 3.07E-02 |

Note: Beta (CNV-Gene), estimated effect size of CNV on gene expression; Beta (CNV-AOD), estimated effect size of CNV on Age of death. P.adj, adjusted P-value calculated based on linear regression by Matrix eQTL and adjusted by FDR. P.adj\_cit, the adjusted P-value from the Causal Inference Test with FDR correction. P.adj\_cit < 0.05.

We further did the functional enrichment analysis for the seven AD longevity-causal genes by using DAVID database (Dennis *et al*. 2003) (**Figure 6**, and **Methods 4.9**). The AD longevity-causal genes were significantly enriched in the plasminogen activation (Enrichment score = 843.9, P = 1.89E-3, FDR = 1.89E-2) and chemotaxis pathways, which involved two genes (*PLAU* and *PLGRKT*). *PLAU* encodes the urokinase-type plasminogen activator (uPA) controlling the key step in the plasminogen activation system (PAS) to convert the plasminogen to plasmin (Mahmood *et al*. 2018). Extensively studies had shown many human diseases controlled by dysfunctional PAS (Fay *et al*. 2007; Kumar *et al*. 2022; Buckley *et al*. 2019; Baker & Strickland 2020). As a plasminogen receptor, *PLGRKT* is a major regulator to promote cell surface plasminogen activation and is highly colocalized with *PLAU* (Andronicos *et al*. 2010). Noted that the expression level of both *PLGRKT* and *PLAU* are positively correlated to AOD and regulated by the copy number dosage of DEL5006 (**Figure 7**), which further indicates the importance of plasminogen activation in longevity.

**Figure 6.**
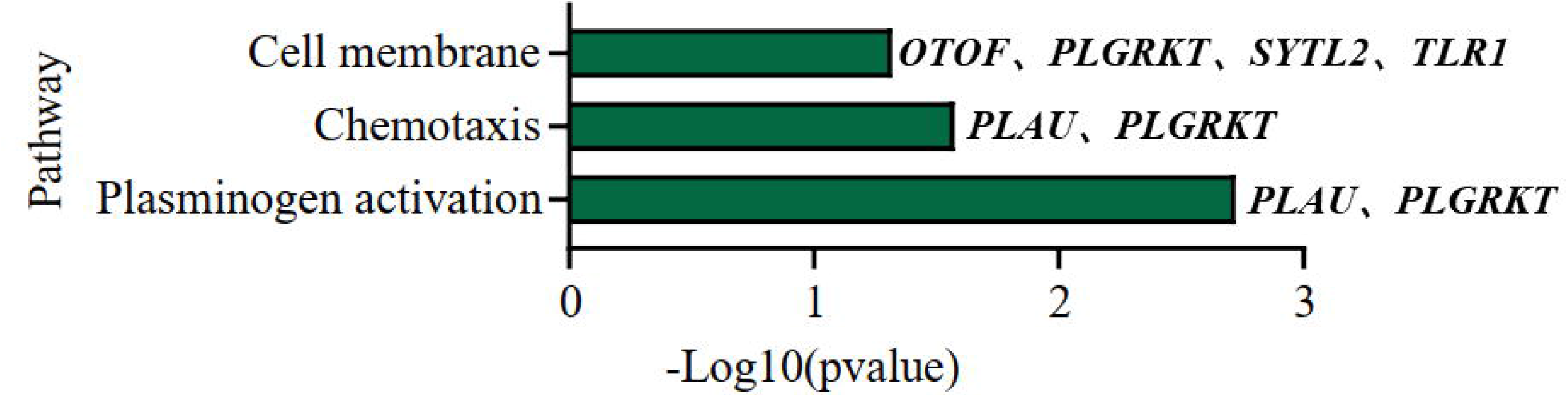
Pathway enrichment analysis of AD longevity-causal genes. The pathways of AD longevity-correlated genes obtained from DAVID. P-value < 0.05, Fisher’s Exact test.

**Figure 7.**
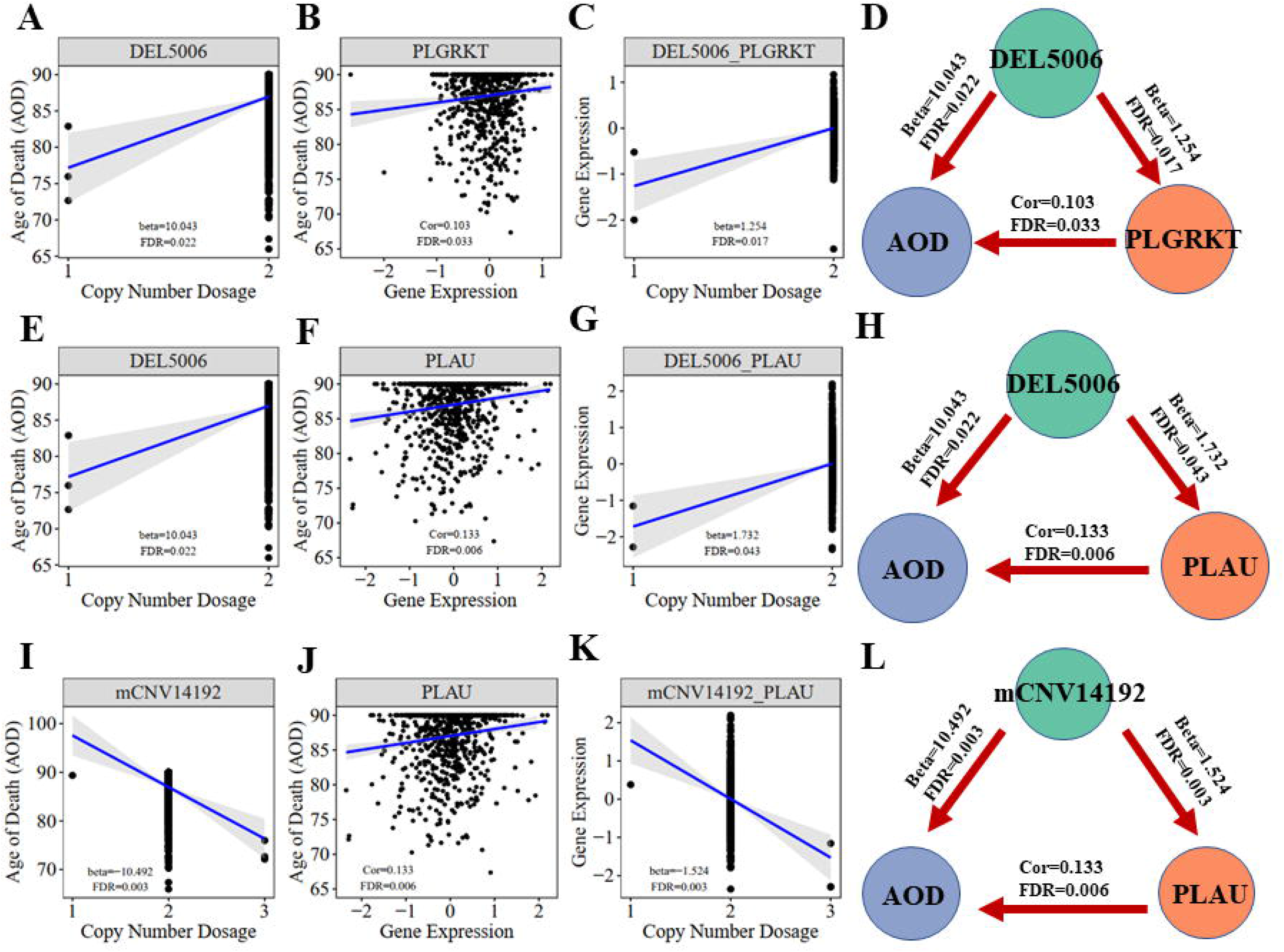
Correlation visualization of DEL5006, mCNV14192, PLGKRT, PLAU, and AOD. (A) Correlation between copy number dosage of DEL5006 and AOD. (B) Correlation between copy number dosage of DEL5006 and PLGRKT gene expression. (C) Correlation between PLGRKT expression and AOD. (D) Causality of DEL5006, PLGRKT, and AOD. (E) Correlation between copy number dosage of DEL5006 and AOD. (F) Correlation between copy number dosage of DEL5006 and PLAU gene expression. (G) Correlation between PLAU expression and AOD. (H) Causality of mCNV14192, PLAU, and AOD. (I) Correlation between copy number dosage of mCNV14192and AOD. (J) Correlation between copy number dosage of mCNV14192and PLAU gene expression. (K) Correlation between PLAU expression and AOD. (L) Causality of mCNV14192, PLAU, and AOD. Beta: estimated effect size of CNV on AOD and gene expression; Cor: spearman coefficient; False Discovery Rate (FDR) < 0.05.

**Figure 8.**
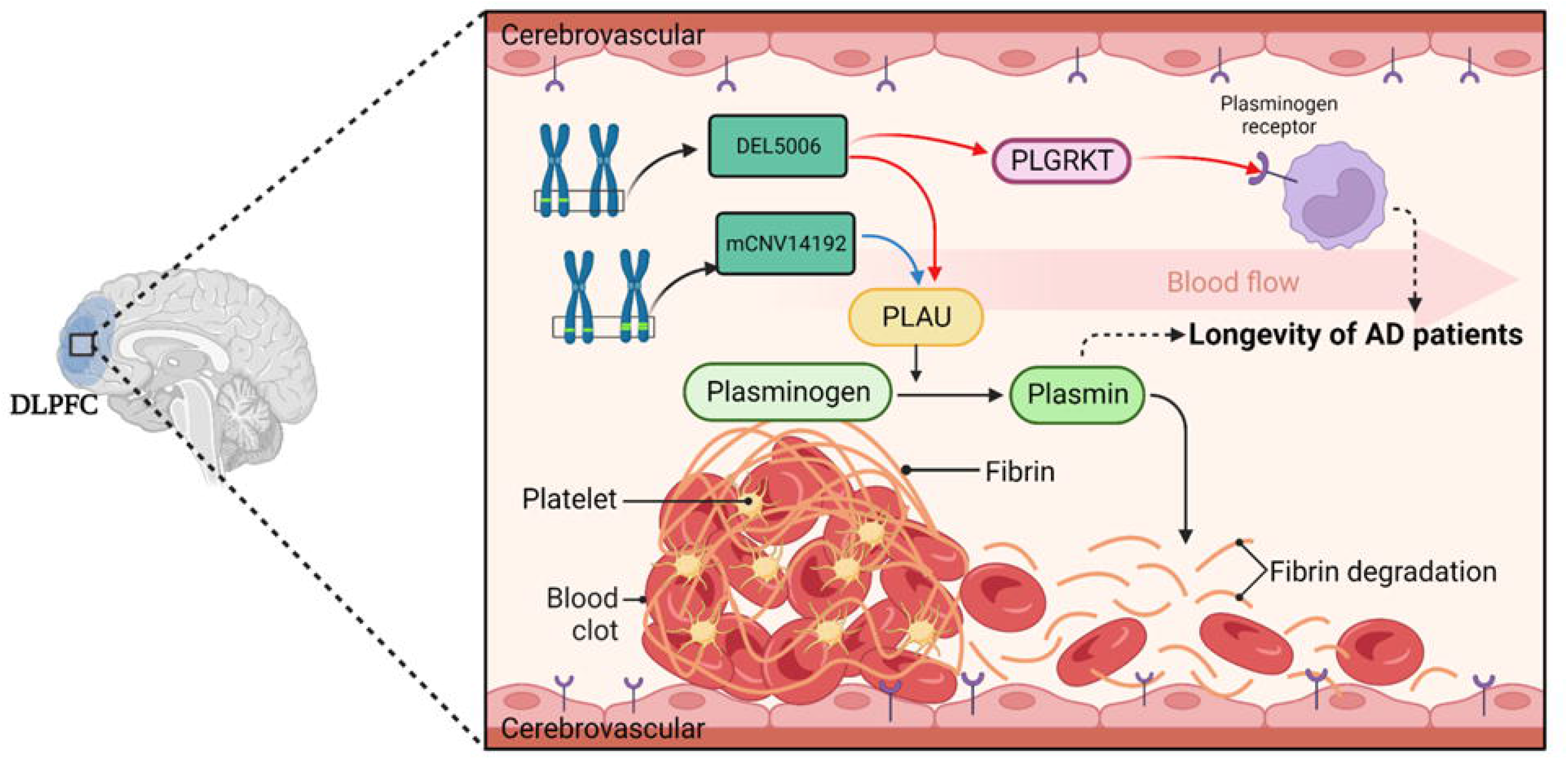
The hypothesis of DEL5006-PLAU/PLGRKT-AOD. The deletion at the DEL5006 loci resulted in the downregulation of *PLGRKT* encoding the plasminogen receptor and *PLAU* encoding the urokinase-type plasminogen activator (uPA), while the duplication at mCNV14192 loci also downregulated *PLAU* expression which caused the inhibition of the conversion from plasminogen to plasmin. This inhibition, in turn, led to the accumulation of fibrin in the cerebrovascular and impaired the longevity of AD patients. The picture was created by BioRender.

## 3 Discussion

The role of CNVs in influencing longevity has been confirmed in various human populations as demonstrated by prior studies (Zhao et al., 2018, Nygaard et al., 2016)(Kuningas *et al*. 2011). Existing research also has shed light on the mechanisms by which genes contribute to longevity (Conneely *et al*. 2012; Melzer *et al*. 2020). Despite these advances, the specific genetic mechanism underpinning longevity of Alzheimer’s Disease patients remains relatively unexplored.

In this study, we have taken a comprehensive approach by integrating CNV profiles, transcriptomic profiles, and meta-information of AD patients based on the ROSMAP cohort. Our construction of a CNV-gene-Age of Death causality network has yielded significant insights. We have successfully identified three critical CNVs - namely, DEL5006, mCNV14192, and DUP42180 - along with seven causal genes correlated with AD longevity. These genes include *PLGRKT*, *TLR1*, *PLAU*, *CALB2*, *SYTL2*, *OTOF*, and *NT5DC1*. The impact of these CNVs and genes on the longevity of AD patients appears to be independent of the severity of their AD status. This discovery provides new perspectives on the genetic influence on longevity in AD patients and opens up fresh avenues for potential therapeutic interventions. Further studies will be beneficial to validate these findings and uncover the precise mechanisms by which these CNVs and genes influence longevity in AD patients.

Our study has led us to propose several intriguing potential hypotheses for further exploration. The role of the plasminogen activation system (PAS), implicated by the identification of the gene *PLAU* and *PLGRKT* in our study, deserves attention. Plasminogen activation has been related to numerous biological processes including tissue remodeling, wound healing, inflammation, fibrinolysis, extracellular migration, cell signaling, cellular migration and degradation of thrombosis within vascular (Baker & Strickland 2020; Castellino & Ploplis). Accumulation of thrombosis makes elders prone to stroke (Kalaria 2000) and may have profound implications for their longevity. The first hypothesis we propose is that CNVs regulate PAS and finally affect cerebrovascular fibrinolysis. The deletion at the DEL5006 loci leads to the downregulation of *PLGRKT* and *PLAU* expression, while the copy number increase at the mCNV14192 loci down-regulates the *PLAU* expression. This downregulation may result in a reduction in plasminogen and plasmin levels, subsequently impairing fibrin degradation within the cerebrovascular system. Such a process could precipitate recurring thrombosis in the brains of Alzheimer’s patients. Consequently, this mechanism may contribute to reduced longevity in DEL5006 carriers (**Figure 7**). Our second hypothesis posits that CNVs modulate the plasmin-independent protective mechanism mediated by uPA-uPAR interaction. It has been observed that uPA treatment can augment the synaptic expression of neuronal cadherin (NCAD) via a uPAR-mediated, plasmin-independent mechanism. Moreover, the formation of NCAD dimers, induced by uPA, has been shown to confer protection to synapses against the harmful influence of soluble Aβ oligomers, as demonstrated in the 5xFAD mouse model (Diaz *et al*.). Another study observed that pharmacologic inhibition or genetic deficiency of plasminogen activator inhibitor-1 (PAI-1) was protective against senescence in mouse model (Eren *et al*. 2014), which further supports our hypothesis of plasminogen activation and longevity. But the exact way the identified CNVs regulate the expression of target genes and how target genes affect the longevity of AD patients remains elusive. Further research is required to understand how these CNVs exert their influence and contribute to AD patient longevity. Moving forward, future work should involve rigorous experimental validation of these hypotheses. In-depth molecular and cellular studies could provide insight into the precise mechanisms underlying these associations.

Our study has several notable limitations. Firstly, the findings, while statistically significant and derived from rigorous analytical methods, lack experimental validations. These findings warrant further verification through experimental studies. Secondly, the identified CNVs and genes’ functional implications, and their interactions, necessitate further scrutiny. An additional constraint is that gene expression was measured solely in the DLPFC region. Consequently, it remains unclear whether the identified causality network is applicable to other brain regions— an aspect which merits further exploration. Furthermore, the population frequency of the AOD-CNVs we identified is low. The correlation coefficient between the genes and AOD is also quite small, implying that there are likely a plethora of other genetic risk factors influencing the longevity of AD patients, which are beyond the scope of the candidate CNVs and genes identified in this study. Human longevity is a complex phenotype, affected by a multitude of genes and environmental factors. Thus, we must interpret our findings within the context of this complexity, acknowledging the likely existence of a myriad of other genetic and environmental contributors to Alzheimer’s patient longevity.

In conclusion, while we have identified key genetic elements that might influence AD patient longevity, we acknowledge that the intricacies of these relationships are likely to be far more complex. Our study provides a steppingstone and offers promising directions for future research in the genetic mechanisms influencing longevity in AD patients.

## 4 Method

### 4.1 CNV profiles based on the Whole-genome sequencing data from the ROSMAP AD cohort

The CNV profiles of the 1,127 individuals from the ROSMAP cohort were downloaded from the AD Knowledge Portal (http://doi.org/10.7303/syn26254632) and the CNV generation details were described in previous study (Ming et al., 2022). These individuals were classified into three groups based on their final Clinical Consensus Diagnosis: 477 individuals with Alzheimer’s disease (AD; scores of 4-5), 285 individuals with mild cognitive impairment (MCI; scores of 2-3), and 365 individuals with normal cognition (NL; score of 1).

### 4.2 Transcriptomic data

The transcriptomics data was downloaded from the AMP-AD portal (synapse ID: syn3388564) and processed in details in previous study (Mostafavi *et al*. 2018b). We further excluded the bias of disease status with function lmFit in R package Limma. We treated AD pathological traits, which are cogdx, braaksc and ceradsc, as covariance independently to exclude the bias from the severity of AD status.

### 4.3 Clinical and pathological trait data

The clinical meta information was downloaded from the AMP-AD portal (Synapse ID: syn3157322). We used the information including age at death, race, sex, braak, ceradsc and cogdx score. To facilitate our analysis, we re-coded the ceradsc score as follows: 1 for Normal, 2 for probable, 3 for possible, and 4 for definite AD. In the correlation analysis, we only considered individuals who were North American whites.

### 4.4 Workflow of this study

In this research, we integrated multi-omics data to explore the potential mechanism of longevity of AD patients. As depicted in **Figure 2**, we primarily utilized Matrix eQTL to identify AOD-correlated CNVs and genes, and subsequently identified presumptive aging causal genes that were correlated with both AOD-correlated CNVs and AOD through Spearman correlation analysis. Next, we inputed the presumptive aging causal genes and AOD information into Rcit to construct a causality network, which we analyzed to reveal the underlying mechanisms of aging.

### 4.5 Correlation analysis of copy number dosage and age of death (AOD)

We used the function Matrix_eQTL_engine with a linear model in the R package Matrix eQTL (version 2.3) (Shabalin 2012) for the correlation analysis. We tested the correlation between CNV dosage and AOD based on linear regression model, as the formula:

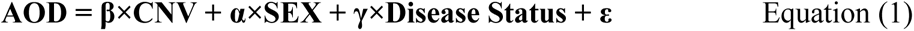

The age of death which were higher than 90 was replaced with 90. The subjects with 0 standard deviations were excluded to make sure the CNV dosage in the whole subjects was different. We used pesudo-code for sex (Female=0, Male=1). There were three AD pathological traits, cogdx, braaksc and ceradsc which were adjusted as well as sex in the generalized linear regression processing and we detected 12, 12, and 11 CNVs correlated with AOD respectively (FDR < 0.05). Finally, we only considered the 10 consensus AOD-correlated CNVs for all three traits.

### 4.6 AOD-correlated CNVs Frequency

To determine the frequency of AOD-correlated CNVs in different groups of the ROSMAP cohort, we calculated the ratio of individuals with AOD-correlated CNVs among the whole cohort, only AD group, and only Normal groups. We further performed a Chi-Squared Test to compare the CNV frequency between AD and Normal groups.

### 4.7 Correlation analysis of CNV dosage and gene expression level

Like the CNV-AOD correlation analysis, Matrix eQTL was also used for the correlation analysis between AOD-correlated CNVs and gene expression. We performed a linear additive model to adjust sex and disease status and got 110 consensus genes whose expression significantly correlated with 10 AOD-correlated CNVs and formed 125 CNV-Gene pairs.

### 4.8 Correlation analysis for the age of death and gene expression

A linear model was applied to exclude bias from disease status of transcriptomics data by using the R package limma’s function lmFit (Ritchie *et al*. 2015). Spearman correction was calculated between log2-transformed expression values output from limma and AOD. Based on cogdx, braaksc, and ceradsc, there were 1294, 2056, and 2774 AOD-correlated genes were identified respectively.

### 4.9 Construct CNV-Gene-AOD correlation network

By intersecting 1294, 2056, 2774 AOD-correlated genes and 110 AOD-CNVs correlated genes, we finally focus on 14, 17, and 21 genes which were both correlated with AOD and AOD-CNVs while using braaksc, cogdx and ceradsc. We used the 17 genes under cogdx forming 23 CNV-Gene-AOD pairs to construct CNV-Gene-AOD correlation network. The correlation pairs were defined as the edges to link their responding nodes. Then Cytoscape (3.9.1) (Shannon *et al*. 2003) was used for network visualization.

### 4.10 Construct CNV-Gene-AOD causality network

The CNV-gene-AOD causality network was built by combining significantly correlated CNV-AOD, CNV-gene, and gene-AOD pairs from the ROSMAP cohort. We compared the probability of two scenarios:

Scenario 1: CNV-Gene-AOD, indicating that CNV contributed to AOD via gene expression change.

Scenario 2: CNV-AOD-Gene, indicating that aging causes the variation of gene expression.

To estimate the likelihood of the two scenarios, we utilized the Causal Inference Test (CIT) (Millstein *et al*. 2016). We choose the scenario with more significant p-value as the more likely scenario. Eventually, we identified three AOD-correlated CNVs that regulate seven genes and affect AOD, which were used to construct the causality network. The correlation pairs between these genes and CNVs were defined as edges that linked their corresponding nodes in the network. We utilized Cytoscape (version 3.9.1) to visualize the network. In the network diagram, pink, yellow, and green circles represent CNVs, genes, and age of death, respectively. The star symbol indicates the gene that is enriched for the chemotaxis pathway, while the purple asterisk represents the AD-risk genes. The red and blue edges represent positive and negative correlations, respectively. In the causality network, arrows are used to indicate the direction of causality.

### 4.11 Functional enrichment

Perform pathways enrichment analysis on seven AD longevity-causal genes using DAVID, with Fisher’s Exact test, as the following equation:

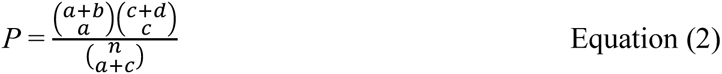

"a" is the genes we input that are mapped to the pathway, "b" is the genes mapped to the pathway based on the whole genome level, "c" is the gene we input that are not mapped to the pathway, "d" is the genes that are not mapped to the pathway on the whole genome level, and "n" represent the total number of genes in the genome (i.e., n = a + b + c + d).

The background was defined as the 18,364 genes detected in the transcriptomics data.

## Acknowledgement

This work was supported by Dr. Chen Ming’s START-UP Research Grant from the University of Macau (SRG2023-00003-FHS). This work was performed in part at the High Performance Computing Cluster (HPCC) which is supported by Information and Communication Technology Office (ICTO) of the University of Macau.

The results published here are in whole or in part based on data obtained from the AD Knowledge Portal (https://adknowledgeportal.org). The data available in the AD Knowledge Portal would not be possible without the participation of research volunteers and the contribution of data by collaborating researchers. Study data were provided by the Rush Alzheimer’s Disease Center, Rush University Medical Center, Chicago. Data collection was supported through funding by NIA grants P30AG10161 (ROS), R01AG15819 (ROSMAP; genomics and RNAseq), R01AG17917 (MAP), R01AG30146, R01AG36836 (RNAseq), U01AG32984 (genomic and whole exome sequencing), U01AG61356 (whole genome sequencing, targeted proteomics, ROSMAP AMP-AD), the Illinois Department of Public Health (ROSMAP), and the Translational Genomics Research Institute (genomic). Additional phenotypic data can be requested at www.radc.rush.edu.

## Reference

C. Reitz, M.A. Pericak-Vance, T. Foroud, and R. Mayeux, A global view of the genetic basis of Alzheimer disease. Nat Rev Neurol 19 (2023) 261–277.doi:10.1038/s41582-023-00789-z

Alzheimer’s Disease International (ADI), Dementia statistics, Available at: https://www.alzint.org/about/dementia-facts-figures/dementia-statistics/ [Accessed Nov 1, 2022],

L.M. Bekris, C.E. Yu, T.D. Bird, and D.W. Tsuang, Genetics of Alzheimer disease. J Geriatr Psychiatry Neurol 23 (2010) 213–27.doi:10.1177/0891988710383571

L. Bertram, and R.E. Tanzi, The genetic epidemiology of neurodegenerative disease. J Clin Invest 115 (2005) 1449–57.doi:10.1172/JCI24761

M.A. DeTure, and D.W. Dickson, The neuropathological diagnosis of Alzheimer’s disease. Molecular Neurodegeneration 14 (2019) 32.doi:10.1186/s13024-019-0333-5

Alzheimer’s Association, Alzheimer’s Disease Facts and Figures, Available at: https://www.alz.org/alzheimers-dementia/facts-figures [Accessed Nov 1, 2022],

R.E. Marioni, S.E. Harris, Q. Zhang, A.F. McRae, S.P. Hagenaars, W.D. Hill, G. Davies, C.W. Ritchie, C.R. Gale, J.M. Starr, A.M. Goate, D.J. Porteous, J. Yang, K.L. Evans, I.J. Deary, N.R. Wray, and P.M. Visscher, GWAS on family history of Alzheimer’s disease. Transl Psychiatry 8 (2018) 99.doi:10.1038/s41398-018-0150-6

J.-C. Lambert, C.A. Ibrahim-Verbaas, D. Harold, A.C. Naj, R. Sims, C. Bellenguez, G. Jun, A.L. DeStefano, J.C. Bis, and G.W. Beecham, Meta-analysis of 74,046 individuals identifies 11 new susceptibility loci for Alzheimer’s disease. Nature genetics 45 (2013) 1452–1458

B.W. Kunkle, B. Grenier-Boley, R. Sims, J.C. Bis, V. Damotte, A.C. Naj, A. Boland, M. Vronskaya, S.J. Van Der Lee, and A. Amlie-Wolf, Genetic meta-analysis of diagnosed Alzheimer’s disease identifies new risk loci and implicates Aβ, tau, immunity and lipid processing. Nature genetics 51 (2019) 414–430

I.E. Jansen, J.E. Savage, K. Watanabe, J. Bryois, D.M. Williams, S. Steinberg, J. Sealock, I.K. Karlsson, S. Hägg, and L. Athanasiu, Genome-wide meta-analysis identifies new loci and functional pathways influencing Alzheimer’s disease risk. Nature genetics 51 (2019) 404–413

D.P. Wightman, I.E. Jansen, J.E. Savage, A.A. Shadrin, S. Bahrami, D. Holland, A. Rongve, S. Børte, B.S. Winsvold, and O.K. Drange, A genome-wide association study with 1,126,563 individuals identifies new risk loci for Alzheimer’s disease. Nature genetics 53 (2021) 1276–1282

C. Bellenguez, F. Küçükali, I.E. Jansen, L. Kleineidam, S. Moreno-Grau, N. Amin, A.C. Naj, R. Campos-Martin, B. Grenier-Boley, and V. Andrade, New insights into the genetic etiology of Alzheimer’s disease and related dementias. Nature genetics 54 (2022) 412–436

A.M. Herskind, M. McGue, N.V. Holm, T.I.A. Sørensen, B. Harvald, and J.W. Vaupel, The heritability of human longevity: A population-based study of 2872 Danish twin pairs born 1870–1900. Human Genetics 97 (1996) 319–323.doi:10.1007/BF02185763

K.N. Conneely, B.C. Capell, M.R. Erdos, P. Sebastiani, N. Solovieff, A.J. Swift, C.T. Baldwin, T. Budagov, N. Barzilai, and G. Atzmon, Human longevity and common variations in the LMNA gene: a meta-analysis. Aging cell 11 (2012) 475–481

D. Melzer, L.C. Pilling, and L. Ferrucci, The genetics of human ageing. Nature Reviews Genetics 21 (2020) 88–101

M. Nygaard, B. Debrabant, Q. Tan, J. Deelen, K. Andersen-Ranberg, A.J. de Craen, M. Beekman, B. Jeune, P.E. Slagboom, K. Christensen, and L. Christiansen, Copy number variation associates with mortality in long-lived individuals: a genome-wide assessment. Aging Cell 15 (2016) 49–55.doi:10.1111/acel.12407

M. Kuningas, K. Estrada, Y.-H. Hsu, K. Nandakumar, A.G. Uitterlinden, K.L. Lunetta, C.M. van Duijn, D. Karasik, A. Hofman, J. Murabito, F. Rivadeneira, D.P. Kiel, and H. Tiemeier, Large common deletions associate with mortality at old age. Hum Mol Genet 20 (2011) 4290–4296.doi:10.1093/hmg/ddr340

C. Ming, M. Wang, Q. Wang, R. Neff, E. Wang, Q. Shen, J.S. Reddy, X. Wang, M. Allen, N. Ertekin-Taner, P.L. De Jager, D.A. Bennett, V. Haroutunian, E. Schadt, and B. Zhang, Whole genome sequencing-based copy number variations reveal novel pathways and targets in Alzheimer’s disease. Alzheimers Dement 18 (2022) 1846–1867.doi:10.1002/alz.12507

P.L. De Jager, Y. Ma, C. McCabe, J. Xu, B.N. Vardarajan, D. Felsky, H.U. Klein, C.C. White, M.A. Peters, B. Lodgson, P. Nejad, A. Tang, L.M. Mangravite, L. Yu, C. Gaiteri, S. Mostafavi, J.A. Schneider, and D.A. Bennett, A multi-omic atlas of the human frontal cortex for aging and Alzheimer’s disease research. Sci Data 5 (2018) 180142.doi:10.1038/sdata.2018.142

S. Mostafavi, C. Gaiteri, S.E. Sullivan, C.C. White, S. Tasaki, J. Xu, M. Taga, H.-U. Klein, E. Patrick, V. Komashko, C. McCabe, R. Smith, E.M. Bradshaw, D.E. Root, A. Regev, L. Yu, L.B. Chibnik, J.A. Schneider, T.L. Young-Pearse, D.A. Bennett, and P.L. De Jager, A molecular network of the aging human brain provides insights into the pathology and cognitive decline of Alzheimer’s disease. Nature Neuroscience 21 (2018a) 811–819.doi:10.1038/s41593-018-0154-9

L.B. Chibnik, C.C. White, S. Mukherjee, T. Raj, L. Yu, E.B. Larson, T.J. Montine, C.D. Keene, J. Sonnen, J.A. Schneider, P.K. Crane, J.M. Shulman, D.A. Bennett, and P.L. De Jager, Susceptibility to neurofibrillary tangles: role of the PTPRD locus and limited pleiotropy with other neuropathologies. Mol Psychiatr 23 (2018) 1521–1529.doi:10.1038/mp.2017.20

A.J. Lee, Y. Ma, L. Yu, R.J. Dawe, C. McCabe, K. Arfanakis, R. Mayeux, D.A. Bennett, H.U. Klein, and P.L. De Jager, Multi-region brain transcriptomes uncover two subtypes of aging individuals with differences in Alzheimer risk and the impact of APOEε4. bioRxiv (2023).doi:10.1101/2023.01.25.524961

S. Chen, L.C. Francioli, J.K. Goodrich, R.L. Collins, M. Kanai, Q. Wang, J. Alföldi, N.A. Watts, C. Vittal, L.D. Gauthier, T. Poterba, M.W. Wilson, Y. Tarasova, W. Phu, M.T. Yohannes, Z. Koenig, Y. Farjoun, E. Banks, S. Donnelly, S. Gabriel, N. Gupta, S. Ferriera, C. Tolonen, S. Novod, L. Bergelson, D. Roazen, V. Ruano-Rubio, M. Covarrubias, C. Llanwarne, N. Petrillo, G. Wade, T. Jeandet, R. Munshi, K. Tibbetts, g.P. Consortium, A. O’Donnell-Luria, M. Solomonson, C. Seed, A.R. Martin, M.E. Talkowski, H.L. Rehm, M.J. Daly, G. Tiao, B.M. Neale, D.G. MacArthur, and K.J. Karczewski, A genome-wide mutational constraint map quantified from variation in 76,156 human genomes. bioRxiv (2022) 2022.03.20.485034.doi:10.1101/2022.03.20.485034

A.A. Shabalin, Matrix eQTL: ultra fast eQTL analysis via large matrix operations. Bioinformatics 28 (2012) 1353–8.doi:10.1093/bioinformatics/bts163

M.E. Ritchie, B. Phipson, D. Wu, Y. Hu, C.W. Law, W. Shi, and G.K. Smyth, limma powers differential expression analyses for RNA-sequencing and microarray studies. Nucleic acids research 43 (2015) e47–e47

J. Millstein, G.K. Chen, and C.V. Breton, cit: hypothesis testing software for mediation analysis in genomic applications. Bioinformatics 32 (2016) 2364–2365.doi:10.1093/bioinformatics/btw135

N. Ertekin-Taner, J. Ronald, L. Feuk, J. Prince, M. Tucker, L. Younkin, M. Hella, S. Jain, A. Hackett, L. Scanlin, J. Kelly, M. Kihiko-Ehman, M. Neltner, L. Hersh, M. Kindy, W. Markesbery, M. Hutton, M. de Andrade, R.C. Petersen, N. Graff-Radford, S. Estus, A.J. Brookes, and S.G. Younkin, Elevated amyloid beta protein (Abeta42) and late onset Alzheimer’s disease are associated with single nucleotide polymorphisms in the urokinase-type plasminogen activator gene. Hum Mol Genet 14 (2005) 447–60.doi:10.1093/hmg/ddi041

S. Liu, Y. Liu, W. Hao, L. Wolf, A.J. Kiliaan, B. Penke, C.E. Rübe, J. Walter, M.T. Heneka, and T. Hartmann, TLR2 is a primary receptor for Alzheimer’s amyloid β peptide to trigger neuroinflammatory activation. The Journal of Immunology 188 (2012) 1098–1107

R. Miskin, and T. Masos, Transgenic mice overexpressing urokinase-type plasminogen activator in the brain exhibit reduced food consumption, body weight and size, and increased longevity. The Journals of Gerontology Series A: Biological Sciences and Medical Sciences 52 (1997) B118–B124

G. Dennis, B.T. Sherman, D.A. Hosack, J. Yang, W. Gao, H.C. Lane, and R.A. Lempicki, DAVID: database for annotation, visualization, and integrated discovery. Genome biology 4 (2003) 1-11

N. Mahmood, C. Mihalcioiu, and S.A. Rabbani, Multifaceted role of the urokinase-type plasminogen activator (uPA) and its receptor (uPAR): diagnostic, prognostic, and therapeutic applications. Frontiers in oncology 8 (2018) 24

W.P. Fay, N. Garg, and M. Sunkar, Vascular functions of the plasminogen activation system. Arteriosclerosis, Thrombosis, and Vascular Biology 27 (2007) 1231–1237

A.A. Kumar, B.J. Buckley, and M. Ranson, The urokinase plasminogen activation system in pancreatic cancer: Prospective diagnostic and therapeutic targets. Biomolecules 12 (2022) 152

B.J. Buckley, U. Ali, M.J. Kelso, and M. Ranson, The urokinase plasminogen activation system in rheumatoid arthritis: pathophysiological roles and prospective therapeutic targets. Current Drug Targets 20 (2019) 970–981

S.K. Baker, and S. Strickland, A critical role for plasminogen in inflammation. J Exp Med 217 (2020).doi:10.1084/jem.20191865

N.M. Andronicos, E.I. Chen, N. Baik, H. Bai, C.M. Parmer, W.B. Kiosses, M.P. Kamps, J.R. Yates III, R.J. Parmer, and L.A. Miles, Proteomics-based discovery of a novel, structurally unique, and developmentally regulated plasminogen receptor, Plg-RKT, a major regulator of cell surface plasminogen activation. Blood, The Journal of the American Society of Hematology 115 (2010) 1319–1330

F.J. Castellino, and V.A. Ploplis, Structure and function of the plasminogen/plasmin system.

R.N. Kalaria, The role of cerebral ischemia in Alzheimer’s disease. Neurobiol Aging 21 (2000) 321–330.doi:https://doi.org/10.1016/S0197-4580(00)00125-1

A.A.-O. Diaz, P. Merino, J.D. Guo, M.A. Yepes, P. McCann, T. Katta, E.M. Tong, E. Torre, S.A.-O. Rangaraju, and M.A.-O. Yepes, Urokinase-Type Plasminogen Activator Protects Cerebral Cortical Neurons from Soluble Aβ-Induced Synaptic Damage.

M. Eren, A.E. Boe, E.A. Klyachko, and D.E. Vaughan, Role of plasminogen activator inhibitor-1 in senescence and aging. Semin Thromb Hemost 40 (2014) 645–51.doi:10.1055/s-0034-1387883

S. Mostafavi, C. Gaiteri, S.E. Sullivan, C.C. White, S. Tasaki, J. Xu, M. Taga, H.-U. Klein, E. Patrick, and V. Komashko, A molecular network of the aging human brain provides insights into the pathology and cognitive decline of Alzheimer’s disease. Nature neuroscience 21 (2018b) 811–819

P. Shannon, A. Markiel, O. Ozier, N.S. Baliga, J.T. Wang, D. Ramage, N. Amin, B. Schwikowski, and T. Ideker, Cytoscape: a software environment for integrated models of biomolecular interaction networks. Genome research 13 (2003) 2498–2504

