## Supplementary Figures for "Effects of Copy Number Variations on Longevity in Late-Onset Alzheimer’s Disease Patients: Insights from a Causality Network Analysis"

### Supplementary Figure

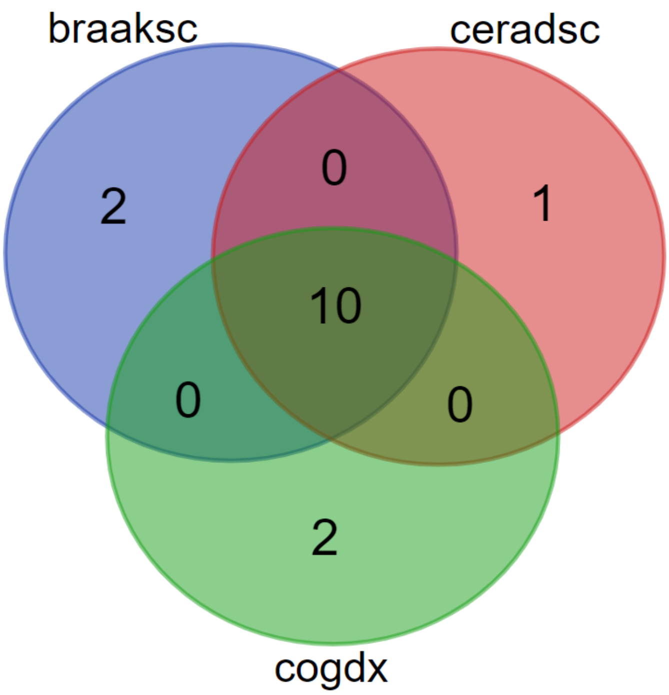


**Figure S1. The Venn plot showing the intersection of AOD-correlated CNVs after adjusting different AD pathologic traits, i.e., braaksc, ceradsc and cogdx, independently.**

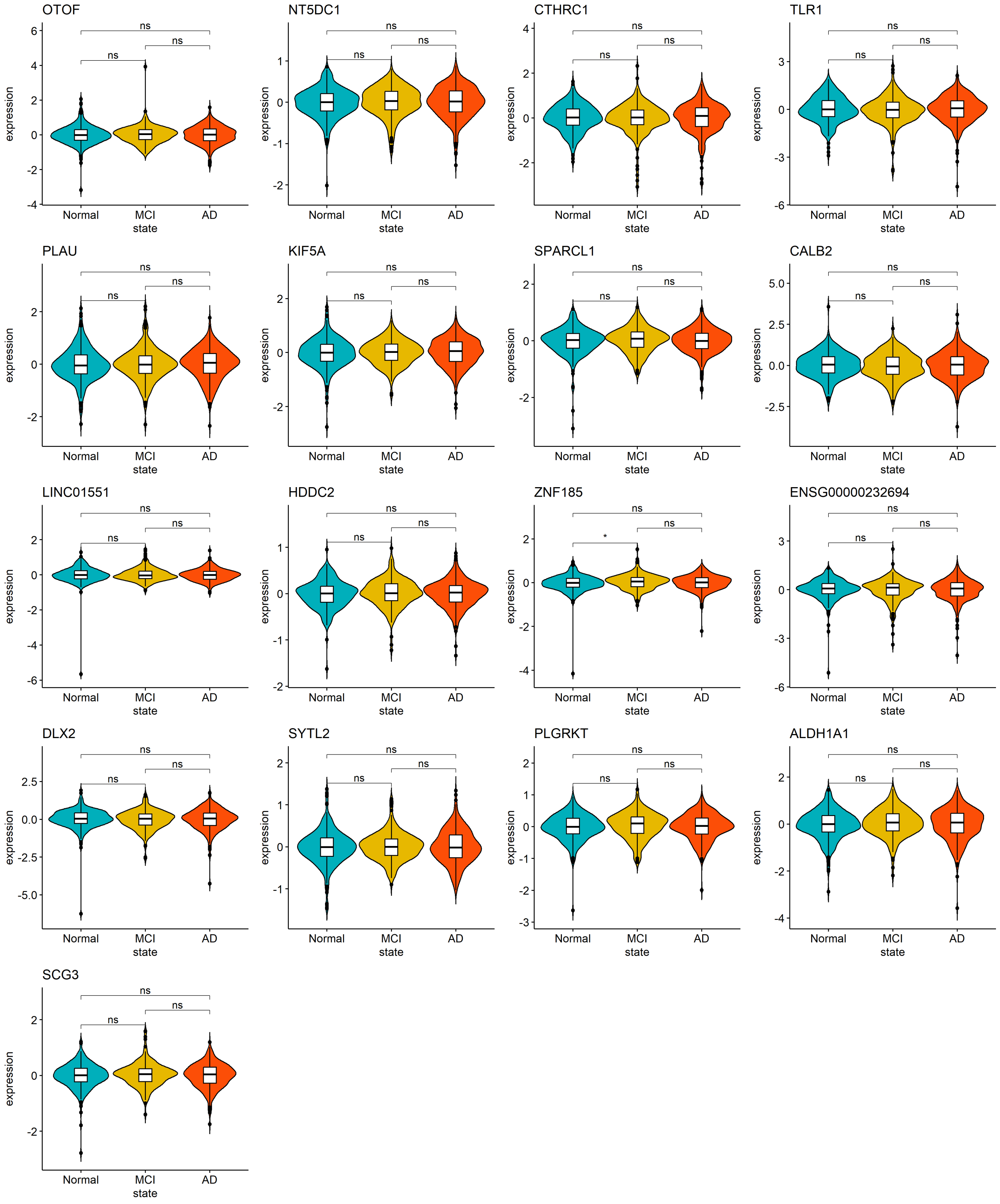


**Figure S2. The violin plot showing the expression level of the presumptive AD longevity-correlated genes after adjusting disease status (cogdx). There is no significant disease status effect on the gene expression level after the co-variance adjustment. The gene expression level is comparable among groups with different disease status.**
